# Learning source weights for target-position estimation: comparing fixed weights and an input-dependent rule

**DOI:** 10.64898/2026.06.08.730995

**Authors:** Jun Kobayashi

## Abstract

Estimating an object’s location requires combining uncertain measurements. Weights derived from an earlier calibration may become inappropriate when measurement errors change. We examined whether learning from noisy reference measurements improves position estimates, and whether weights computed from current inputs improve further than one learned set of weights used throughout evaluation. In simulations, we generated noisy position measurements of a target that remained fixed within each trial. Each two-input method combined the current measurement with the mean of three initial measurements. Measurement-accuracy reductions and added measurement offsets applied only after initialization. The estimator received no current or previous true target position. We repeated coordinate calibration, training, validation, and independent evaluation 100 times in each of four error conditions. Recalibration and fixed-weight learning explained most of the improvement over earlier supplied weights under reduced measurement accuracy and temporal correlation. With occsional large measurement errors, input-dependent weighting reduced mean squared error by 4.05% relative to learned fixed weights and assigned lower mean weights to measurements with added offsets. Its additional benefit was below 0.5% under independent errors and reduced current accuracy, while fixed weights had slightly lower mean error under temporal correlation. Methods using longer observation histories performed better in several conditions. Sensitivity analysis showed that shared coordinate bias and correlated reference errors limit conclusions about physical accuracy. The results show benefits and limits of recalibration, input-dependent weighting, and observation history within these measurement models.

## 1 Introduction

Estimating an object’s position requires combining uncertain information. Perceptual studies have established methods for measuring the influence of individual cues and relating it to their reliability (Young et al, 1993; Landy et al, 1995; Ernst and Banks, 2002). Position estimates used in reaching likewise depend on uncertain information (Körding and Wolpert, 2004; Sober and Sabes, 2005). Here we examine a computational question: can learning improve position estimates when the uncertainty values used to combine inputs no longer adequately describe their errors?

Inverse-variance weighting gives greater influence to inputs with smaller specified error variances. In this study, the error estimates came from an earlier calibration; we call the resulting weights supplied weights. Incorrect variance specifications can therefore give an unreliable input too much influence. Human perceptual judgments can also be modeled using uncertainty estimates that differ from measured sensory variability (Hong et al, 2025). Moreover, errors shared by inputs are not described by their individual variances alone (Oruç et al, 2003). These issues motivate comparing weights calculated from calibration data with weights learned from subsequent measurements and feedback.

Feedback can change visual cue weights (Jacobs and Fine, 1999; van Beers et al, 2011), and sensory uncertainty can be learned from experience (Sato and Körding, 2014; Beierholm et al, 2020). We first ask how much an earlier calibration can be improved by re-estimating input errors or learning one set of weights from subsequent reference measurements. We then ask whether a rule that computes weights from current inputs improves further. In our two-input model, fixed-weight fitting and covariance-based weighting that includes cross moments yield the same solution for the same training data and constrained squared-error objective; their agreement does not distinguish learning mechanisms.

Occasional large errors make the distinction concrete. Reducing the weight of a contaminated input on every measurement can improve estimates without identifying individual errors. An input-dependent rule can instead use disagreement to reduce that input’s influence selectively. This question relates to robust integration and causal-inference models (Knill, 2007; Körding et al, 2007); disagreement alone, however, does not establish that the inputs arise from different physical sources.

We study a fixed, unknown three-dimensional target using simulated position measurements. A prediction formed from initial measurements is combined with the current measurement. Both inputs come from repeated measurements within the same simulated measurement process, rather than distinct perceptual cues or sensory modalities. The estimator does not receive the target’s true position. Training uses noisy reference measurements rather than exact targets. We compare learning and recalibration under changes in measurement accuracy, occasional large measurement errors, and temporal error correlation. Comparisons with methods that retain all available measurements assess the limits of compressing the history into an initial prediction. We also examine how uncertainty in coordinate calibration and reference measurements limits the conclusions. The study evaluates computational estimation rules, not human learning mechanisms or executed reaches.

## 2 Methods

### 2.1 Estimation task and available information

Each trial contained an unknown target position that remained fixed during 23 position measurements; target positions differed between trials. These were repeated measurements from one simulated position-measurement process, rather than measurements from separate sensory modalities. The mean of the first three measurements initialized a position prediction; the remaining 20 were evaluated. For each coordinate, let *c*_*j*_ denote position measurement *j* and define the initial prediction as 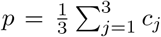. Evaluated time points are indexed by *t* = 1, …, 20, with current measurement *v*_*t*_ = *c*_*t*+3_. Each two-input weighting method estimated position as

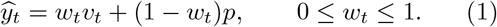

All two-input methods used the same initial prediction, which remained unchanged within a trial. Because the target was stationary, the initial mean provided a prediction of its position at subsequent times. Averaging multiple initial measurements reduces their random error when the errors are independent; temporal correlation changes that reduction. The choice of three initialization measurements was not optimized.

With this common prediction, differences between the two-input estimates reflect how the methods weight the same inputs, rather than differences in how they construct a prediction. We learned either weights shared across evaluation inputs or a rule that computes weights from the current measurement and initial prediction. We did not learn a temporal prediction model, such as an autoregressive model. This comparison concerns weighting with a fixed prediction, rather than the best estimator using all available history. Controls that incorporate later measurements into the history are described in Section 2.3.4. Fig. 1 shows the information used by the methods.

**Fig. 1.**
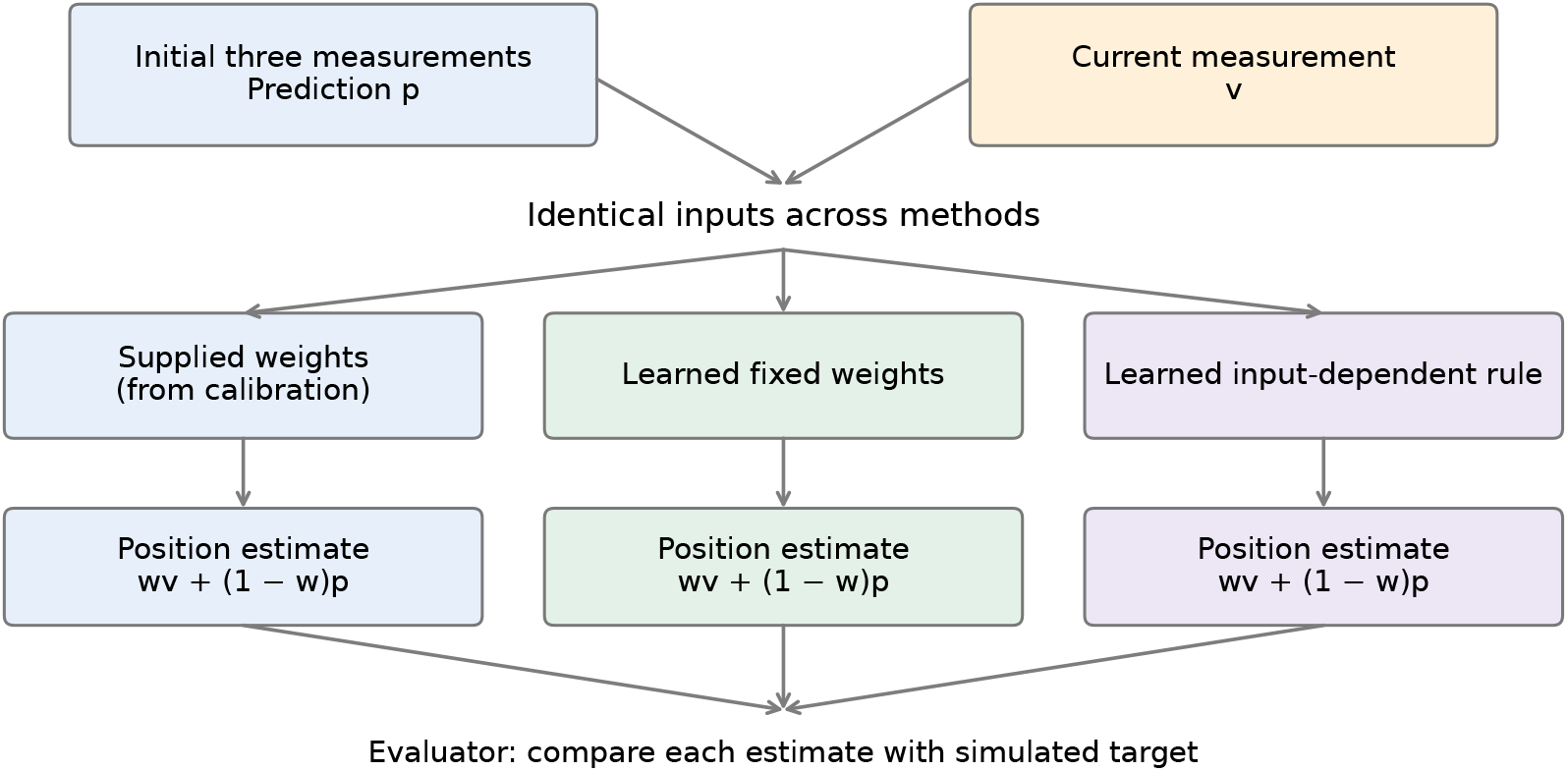
Position estimation from observable inputs. Initial position measurements provide a fixed prediction *p*; a later position measurement provides *vt*. Each weighting method combines these two inputs to produce its own position estimate. The evaluator compares each estimate with the simulated target. Training references and the evaluator target are not inputs to the weighting methods during evaluation.

Offline training used the mean of two noisy reference measurements of the target. The estimator received only position measurements during evaluation; reference measurements, target identity, and true positions were excluded from its input. True positions were used to generate simulated measurements and score estimates. The evaluation did not provide feedback or update any learned parameters.

### 2.2 Measurements and coordinate calibration

Each target coordinate was sampled independently from a uniform distribution within *±*0.05 m of the corresponding coordinate of [ *−*0.175,*−*0.245, 1.405] m, defining a cubic workspace. These values define the simulated workspace, not a measured distribution of human targets. We generated three-dimensional position measurements by adding errors to target coordinates. These readings represent a simplified model of position information that could be obtained from a camera. We did not simulate image formation, position detection, or a tracking algorithm. The primary position measurements had independent zero-mean Gaussian errors with SD 0.01 m per coordinate before the condition-specific changes described below. Reference instruments A and B and a separate coordinate standard also had zero-mean Gaussian measurement errors, with per-coordinate SDs 0.002 m for each reference and 0.0005 m for the standard. In the primary setting, measurement errors were independent across instruments, coordinates, and targets. Condition-specific temporal dependence and shared reference errors are described below. These are specified simulation hypotheses, not empirically validated sensor accuracies.

The primary sensor and reference instruments had coordinate offsets of [0.030, *−*0.020, 0.010], [0.028, *−*0.021, 0.013], and [0.033, *−*0.018, 0.008] m, respectively. We estimated each instrument’s translation offset from the mean difference between its readings and independent noisy standard readings. The primary sensor supplied its first measurement for this calibration. Corrected readings subtracted the estimated offset. Different targets were used to validate the corrections. Calibration used measured standard readings, not hidden target coordinates.

The primary simulation assumed independent, zero-mean standard and reference errors. Shared errors cannot generally be identified from instrument differences (Joint Committee for Guides in Metrology, 2008). Accordingly, the sensitivity analysis added standard bias and correlated reference errors. Translation calibration does not address scale or rotation errors, and the standard’s systematic uncertainty was not inferred from the position measurements.

### 2.3 Weighting methods

#### 2.3.1 Weights from calibrated error moments

For each coordinate, define the two-input error vector *e* = [*v −y, p −y*]^T^ and its second-moment matrix *M* = E[*ee*^T^]. We estimated *M* from residuals against noisy reference means, rather than observing *e* directly. Its diagonal entries include variance and squared bias; we do not interpret them as centered variances when biases remain.

Let *a* and *b* be corrected reference readings and let 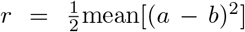. Under independent, unbiased reference errors, *r/*2 estimates the error variance of their mean, including when the two instruments have unequal precision. We subtracted *r/*2 from every entry of the residual second-moment matrix because the same reference error enters both source residuals. Correlated reference errors violate this correction. Finite calibration errors can also leave residual biases; repeating the complete calibration procedure captures variation from those errors without making the correction exact. In the primary setting, the primary sensor’s translation correction has an expected SD of approximately 0.63 mm per coordinate, from the primary-sensor and standard noise averaged over 256 calibration targets. Its error enters both inputs of every target in a run and is not removed by subtracting reference noise.

We call weights calculated from the source-calibration group supplied weights, and weights recalculated from subsequent training data training-recalibrated weights. Supplied-MSE weighting uses the calibrated diagonal entries, which include error variance and squared bias:

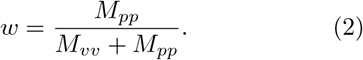

This is the usual inverse-variance form when the inputs are unbiased and uncorrelated. Supplied-covariance weighting also used the cross moment:

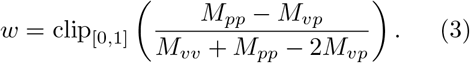

Both weighting methods used the same second-moment matrix, estimated from a separate source-calibration group. We also recalculated these weights from weight-training data, so that recalibration and direct learning could use the same subsequent reference measurements. A further comparison used a separate training matrix for each evaluation time point. Matrices were required to be positive definite; we did not regularize or repair invalid matrices.

#### 2.3.2 Learned fixed weights

For each coordinate, we learned a weight for the current measurement by minimizing training MSE against the noisy reference means. For source differences *d* = *v − p* and training references *z*, the fitted weight was

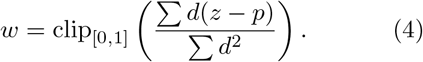

Sums run over training targets and evaluated time points. Using the same training residuals, Eq. 3 reduces algebraically to Eq. 4: subtracting a common reference-noise term from all matrix entries cancels from both numerator and denominator. We retained both implementations as a consistency check, not as distinct estimators. The three learned weights, one for each coordinate, were used unchanged across evaluation targets and time points. Averaging over time within training targets does not make those time points independent trials.

#### 2.3.3 Input-dependent weighting

We learned a separate nine-parameter rule, with three parameters per coordinate:

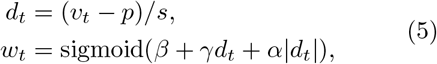

Where 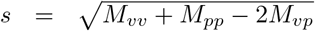 came from source calibration. The intercept represents a constant weight, the signed difference allows asymmetric responses, and its magnitude allows responses to either sign of disagreement. This feature choice was designed for the present comparison; it is not a fitted model of a human learning mechanism. Setting both slopes to zero makes the weight independent of the inputs. The initial intercept reproduced supplied-MSE weighting, with both slopes zero. The calibration matrix, feature scale, and selected parameters were fixed during evaluation.

Training minimized MSE against noisy reference means. Adam (Kingma and Ba, 2015) used learning rate 0.02 for at most 200 updates, with stopping after 20 validation checks without improvement. We then refined the validation-selected parameters with L-BFGS. Each outer step allowed 20 internal iterations with a strong-Wolfe line search; at most 60 outer steps were allowed. Five outer checks without validation improvement stopped this stage. Initial parameters and all subsequent checkpoints were eligible for selection by minimum validation MSE. Training used float64 arithmetic and no minibatching. The L-BFGS objective was divided by the mean squared feature scale to improve numerical conditioning. This fixed positive scaling does not change its minimizer. Optimizer limits were fixed after development experiments, before final evaluation data were generated. No final run reached the L-BFGS limit. Validation stopping does not prove global optimality.

#### 2.3.4 Additional controls using observable history

We evaluated the current measurement alone, the initial prediction alone, and equal weighting. We also evaluated the mean and coordinate-wise median of all position measurements available up to the current time. At the first evaluated time point, history methods and two-input rules use the three initialization measurements and the current measurement. Thereafter, history methods retain intermediate measurements that the two-input rules discard. These comparisons assess the limits of the two-input representation; they do not isolate the effect of the learned weighting procedure. History-based generalized least squares (GLS) combines all measurements available up to the current time, using weights calculated from their error variances and correlations estimated from training data. The calculation used the full error-moment matrix, which also includes bias, and corrected the estimated moments for reference noise. It may use negative weights. If its estimated history matrix was not positive definite, GLS was recorded as unavailable; other valid methods from that run were retained.

### 2.4 Error conditions and independent evaluation

We evaluated four conditions (Table 1). Source calibration used the original independent measurement errors. The changed condition then applied to weight training, validation, and evaluation. Thus, comparisons against supplied weights assess updating an earlier calibration, whereas comparisons against training-recalibrated weights assess the additional benefit of the input-dependent rule. The reduced-current-accuracy and large-error conditions assume that the first three measurements are acquired before accuracy deteriorates or added measurement offsets occur. These initial measurements still contain Gaussian measurement error; they are not exact target positions.

**Table 1.** Measurement errors during weight training, validation, and evaluation. Coordinate calibration and source calibration used independent errors with SD 0.01 m in all conditions. All methods received the same measurements within a run. The target remained fixed in every trial.

| Condition | Measurement errors |
| --- | --- |
| Independent errors | Independent Gaussian errors, SD 0.01 m. |
| Reduced current accuracy | Current-measurement SD increased to 0.03 m after the three initialization measurements. |
| Occasional large measurement errors | Each evaluated measurement independently received an additional vector of length 0.05 m with probability 0.08; direction was uniform on the sphere. Initialization measurements had no added offsets. |
| Temporal correlation | Temporally correlated errors followed a first-order autoregressive process (AR(1)) with coefficient 0.8 through all 23 measurements; marginal SD remained 0.01 m. |

Each condition comprised 100 independent repetitions of the complete procedure. Every repetition generated seven groups of 256 distinct targets: coordinate calibration, coordinate validation, reference-noise calibration, source calibration, weight training, weight validation, and evaluation. No target was shared between groups. Master seeds were generated from a separate seed sequence from development experiments. Separate random streams controlled targets, measurement noise, each reference instrument, coordinate standards, and large measurement errors. Rule parameters were initialized deterministically from each run’s calibration.

The number of runs, sample counts, methods, and optimizer limits were fixed before generating the final evaluation data. Development used 12 repetitions per sensitivity setting with 128 targets per group. Increasing the final group size to 256 provided more measurements for history-moment estimation; 100 complete repetitions characterized variation in calibration and learning.

### 2.5 Outcomes, sensitivity, and verification

We first averaged squared position errors over targets, evaluated time points, and coordinates within each complete run:

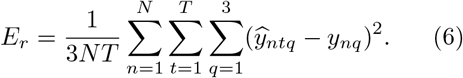

Here *N* = 256 and *T* = 20 in the final experiment. MSE averages the squared errors over the three coordinates, rather than summing them. We then summarized the 100 run-level MSEs in m^2^. Paired differences compared methods within runs; a positive fixed-minus-input-dependent difference favors the learned rule. Percentage differences divided differences between mean MSEs by the named comparator’s mean MSE. We computed run SD and Monte Carlo standard error and report two-sided Student-*t* 95% intervals for the mean of paired differences. These describe numerical uncertainty under the specified simulation, not uncertainty about human performance or biological validity. Intervals were exploratory and unadjusted for multiple comparisons; no equivalence test was performed. Additional post-hoc analyses summarized saved errors at each time point, including the first evaluated measurement (Appendix A); the main outcome remained the full-sequence MSE in Eq. 6.

For large measurement errors, offset indicators available only to the evaluator distinguished measurements selected to receive offsets from other measurements. We compared the mean weight assigned to the current measurement between these groups. These indicators were not an estimator feature, training label, or source of parameter selection. A lower weight on the offset group is descriptive evidence of selective weighting in this simulation, not evidence of causal inference.

Development sensitivity analyses repeated the complete procedure with standard biases of 1, 5, or 10 mm in one coordinate, reference-error correlations of 0.5 or 0.9, reference-B SD multiplied by two or four, and a combined 5-mm bias, 0.5 correlation, and doubled reference-B SD. Each setting and the original measurement setting had 12 independent repetitions per measurement-error condition. Different settings used different seed streams. These settings illustrate possible departures from the measurement assumptions; their frequencies and magnitudes were not estimated from real instruments.

All 400 final runs passed checks for disjoint target groups, exclusion of true positions and evaluation references from fitting, and fixed parameters during evaluation. Changing future measurements left past estimates unchanged; changing evaluator targets affected scoring without changing predictions. A separate NumPy implementation verified predictions, MSEs, coordinate corrections, source moments, and selected validation losses for the final and development experiments. These checks establish numerical reproducibility under the stated model. Raw measurements, fitted parameters, training histories, predictions, and evaluation outputs were retained with SHA-256 manifests. Detailed optimization records, verification counts, and checks are documented in `VERIFICATION.md` in the standalone package’s `scripts/revision observable target/`directory. The package is located at `releases/ observable-v0.2.0/`in the archived repository (see Statements and Declarations).

### 2.6 Use of AI assistance

The author used OpenAI Codex and Anthropic Claude Code to assist with study planning, code development and debugging, simulation and analysis workflows, verification scripts, interpretation of results, figure preparation, and manuscript drafting and revision. The author directed the work and made decisions about the study design and scope of the claims. Numerical checks and independent recalculations are described above. The author retains responsibility for the methods, results, interpretations, and final manuscript.

## 3 Results

### 3.1 Updating an earlier calibration

All 400 final runs completed, with 100 independent repetitions of calibration, training, and evaluation in each measurement-error condition. The input-dependent rule reduced mean MSE relative to supplied-MSE weighting by 57.33% under reduced current accuracy and 9.34% under temporal correlation (Table 2). Learned fixed weights and training-recalibrated weights obtained most of these reductions. In the independent-error condition, the input-dependent rule’s reduction relative to supplied-MSE weighting was 0.36%. These comparisons concern updating an earlier calibration, rather than a comparison with known true variances.

**Table 2.** Mean run-level MSE across 100 complete independent repetitions. All entries are in 10^*−*5^ m^2^. Training-covariance weighting and learned fixed weighting give the same estimator under the shared training data and convex two-input objective. History methods retain all measurements up to and including the current measurement. After the first measurement following the three initialization measurements, history methods use more measurements than the two-input rules.

| Method | Independent errors | Reduced accuracy | Large errors | Temporal correlation |
| --- | --- | --- | --- | --- |
| Supplied MSE | 2.541 | 7.530 | 2.971 | 5.922 |
| Recalibrated MSE | 2.542 | 3.228 | 2.829 | 5.360 |
| Recalibrated covariance / learned fixed | 2.541 | 3.228 | 2.828 | 5.363 |
| Input-dependent | 2.532 | 3.213 | 2.714 | 5.368 |
| History mean | 0.989 | 5.680 | 1.386 | 4.908 |
| History median | 1.379 | 4.995 | 1.505 | 5.302 |
| History-based GLS | 1.040 | 2.604 | 1.392 | 4.777 |

### 3.2 Additional benefit of input-dependent weighting

The input-dependent rule’s largest additional benefit occurred with large measurement errors. Its mean MSE was 2.7140 *×* 10^*−*5^ m^2^, compared with 2.8285 *×* 10^*−*5^ m^2^ for learned fixed weights, a reduction of 4.05%. This direction held in all 100 runs. The paired mean difference was 1.145 *×* 10^*−*6^ m^2^ (95% Monte Carlo interval [1.102, 1.187] *×* 10^*−*6^ m^2^; Table 3; Fig. 2).

**Table 3.** Learned-fixed minus input-dependent MSE across independent runs. Positive differences favor the input-dependent rule. Intervals describe Monte Carlo uncertainty in the mean under the simulated protocol.

| Condition | Mean difference<br>($10^{-6} \text{ m}^2$) | 95% interval<br>($10^{-6} \text{ m}^2$) | Lower MSE<br>with rule |
| --- | --- | --- | --- |
| Independent errors | 0.093 | [0.070, 0.117] | 84/100 |
| Reduced current accuracy | 0.151 | [0.089, 0.212] | 65/100 |
| Occasional large measurement errors | 1.145 | [1.102, 1.187] | 100/100 |
| Temporal correlation | -0.053 | [-0.103, -0.002] | 40/100 |

**Fig. 2.**
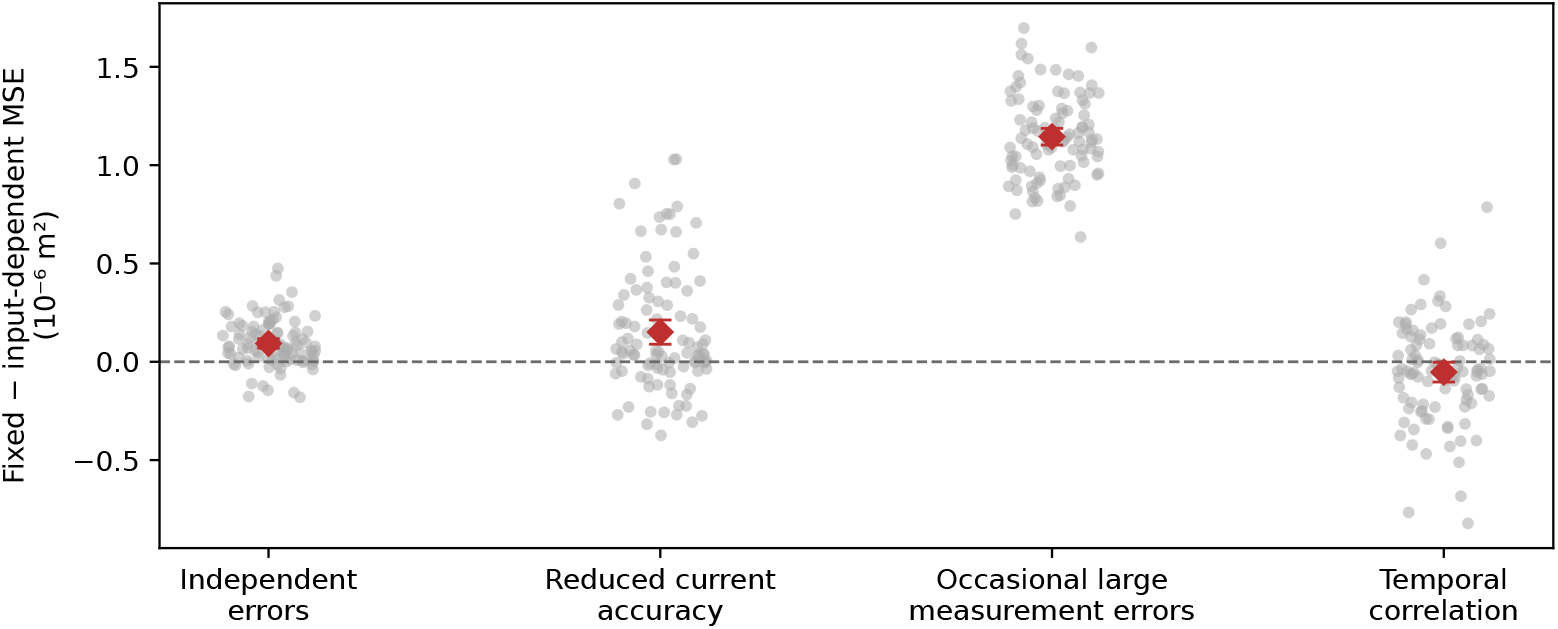
Additional benefit over learned fixed weights. Gray points show paired MSE differences for each complete run; red diamonds and intervals show means and 95% Monte Carlo intervals. The dashed line marks equal MSE. Positive values favor input-dependent weighting.

The corresponding mean reductions relative to learned fixed weights were 0.37% under independent errors and 0.47% under reduced current accuracy. The rule had lower MSE in 84 and 65 runs, respectively. Under temporal correlation, the rule’s mean MSE was 0.098% higher than learned fixed MSE; it had lower error in 40 runs. The absolute differences were small, and the intervals do not establish practical equivalence.

In the large-error condition, mean weight assigned to the current measurement was 0.268 without added offsets and 0.175 on measurements selected to receive offsets. The mean clean-minus-offset difference was 0.093 (95% interval [0.091, 0.096]). The mean within-run relative reduction was 34.7% relative to the clean weight. The difference was positive in all 100 runs. These labels were used only for analysis, not to select the weights.

### 3.3 History controls and measurement sensitivity

Using all available history produced larger benefits than the two-input rule in several settings. History averaging had lower mean MSE than the input-dependent rule in the independent-error, large-error, and temporal-correlation conditions. Under reduced current accuracy, simple history averaging was worse: it gave equal weight to the three initial measurements with SD 0.01 m and the noisier subsequent measurements with SD 0.03 m. History-based GLS used the estimated full error-moment matrix of the available history and had the lowest mean MSE. GLS also had the lowest mean MSE under temporal correlation. Thus, the additional benefit over learned fixed weights does not establish that the rule is preferable to estimators with access to longer observation histories. Appendix A reports every method and the time course of error for selected comparisons.

Development sensitivity analysis completed 432 valid primary comparisons. All 12 independent-error runs and seven large-error runs with fourfold reference-B SD had unavailable history-based GLS estimates; other methods were retained. A standard bias shifted corrected measurements and training references together, leaving that shared coordinate error unobservable to the estimator. The additional mean benefit under large measurement errors was smaller in the 5- and 10-mm standard-bias settings, and small differences between methods changed sign in some other settings (Appendix B). Each setting used different measurement draws, so these are descriptive comparisons rather than paired effects of bias. A shared coordinate shift can also change relative MSE reductions without changing fitted weights. Correlated reference errors also violate the difference-based reference-noise correction. These exploratory results do not justify a general robustness claim across uncertain measurement systems.

## 4 Discussion

### 4.1 What learning added beyond recalibration

We assessed improvement over an earlier uncertainty calibration and the additional benefit of adapting weights to current measurements. Re-estimating input error moments explained most of the benefit over the original supplied weights when measurement accuracy changed. The further benefit of the input-dependent rule was largest for occasional large measurement errors. It was small under independent errors and reduced current accuracy, and its mean MSE was slightly higher under temporal correlation. These results support condition-specific conclusions about the tested rules, rather than a general need for input-dependent weighting.

Learned fixed weights and training-covariance weights were the same solution in this two-input convex estimator. This agreement follows from the shared squared-error objective, not from two independent demonstrations of performance. Direct weight fitting and learning error statistics can therefore give the same estimator when they use the same measurements and model class. Diagonal-only weighting omits cross moments and need not be identical. The supplied weights additionally used earlier calibration data, so improvement over them cannot by itself show that nonlinear weighting was necessary.

With independent Gaussian errors, finite-sample calibration and learning still produce imperfect weights. The learned rule could retain its calibration-based initialization or another validation-selected checkpoint, whereas the fixed fit used the training minimum directly. The small independent-error advantage therefore does not challenge optimal inverse-variance weighting with known independent error statistics or isolate the effect of nonlinear features from model selection. An absence of an advantage in a setting would likewise not establish equivalence.

### 4.2 Disagreement, large measurement errors, and observation history

The input-dependent rule contained a direct magnitude-of-disagreement feature. In the large-error condition, the mean weight assigned to the current measurement was lower when that measurement received an added offset, accompanying lower MSE than learned fixed weights. This result concerns a simulated contamination mechanism and the specific architecture. Disagreement remains ambiguous: a large difference can arise from an error in the current measurement, the initial prediction, or both. Initialization measurements contained Gaussian errors but no added offsets, which made the prediction a comparatively useful reference. Performance with initial large measurement errors or moving targets remains unestablished.

Robust integration and causal-inference models address disagreement in human cue combination (Knill, 2007; Körding et al, 2007). Here the current measurement and prediction were derived from measurements of the same fixed target at different times. No common-cause variable was inferred, and we did not compare causal-inference estimators. Selective weighting in this simulation therefore cannot establish a human outlier-detection or causal-inference mechanism.

The history controls limit the practical interpretation of the two-input comparison. Simple averaging of all available measurements had lower mean MSE in three conditions, including large measurement errors. A history-based GLS estimate had the lowest mean MSE under reduced current accuracy and temporal correlation. After the first evaluated measurement, these methods retain measurements that the two-input rules discard.

At the first evaluated measurement, however, input-dependent weighting had lower MSE than history averaging and history-based GLS in the large-error condition, when all methods used the three initialization measurements and the current measurement (Appendix A). The full-sequence advantage of history methods therefore does not imply that their integration procedure was better at every time point. These comparisons show benefits of disagreement-dependent weighting and accumulating measurements in the tested conditions, without isolating their causal contributions. The time courses in Appendix A show how history averaging improves as measurements accumulate, while the two-input methods retain the same initial prediction. Under independent errors, the expected error of the history mean includes both a noise term decreasing with the number of measurements and a shared translation-calibration error that averaging cannot remove; the other conditions need not follow that independent-noise pattern.

### 4.3 Measurement uncertainty and biological interpretation

Learning cannot remove a common error that neither its inputs nor its reference measurements reveal. A biased coordinate standard shifted the corrected primary measurements and training references together. The learned disagreement weights could remain unchanged while absolute position estimates became biased. Similarly, correlated reference errors are hidden from differences between reference instruments. Improving estimates relative to such references is insufficient to establish improvement in physical accuracy. This is an identifiability limit of the measurement arrangement, not a failure that can necessarily be resolved by longer training.

The simulations make information available to the estimator explicit: observed position coordinates during evaluation and noisy reference measurements during offline learning. The procedure also includes independent repetitions of coordinate calibration and learning. However, the primary sensor, reference instruments, and coordinate standard remain generated measurement models. The work provides a controlled comparison under those models, not empirical validation of a sensing system.

Human studies demonstrate experience-dependent cue weights and learning of sensory uncertainty (Jacobs and Fine, 1999; Sato and Körding, 2014; Beierholm et al, 2020), and sensory weighting during motor planning depends on the estimation problem (Sober and Sabes, 2005). The present results suggest useful computational distinctions for such studies: updated average weights, weighting based on current disagreement, and use of a longer observation history need separate comparisons. We did not measure human performance, identify a biological learning rule, or establish that these measurement-error conditions describe human sensory noise.

Target-estimation differences also do not directly determine reaching endpoints. Testing that link requires a controller, assumptions about feedback delay, and assessment of how muscle redundancy and co-contraction change the response to an estimate (Todorov and Jordan, 2002; Osu et al, 2003). Such an evaluation also requires validated body-state estimates and a muscle controller driven by observable inputs and estimated targets.

### 4.4 Limits and future tests

The target remained fixed, and the primary sensor and reference instruments used a shared coordinate representation after translation correction. Scale, rotation, image formation, occlusion, packet delays, and persistent large measurement errors were not tested in the final study. Error levels were specified rather than measured. The sensitivity analysis varied selected reference assumptions; it does not establish robustness to every measurement failure. Its 12-run summaries also have larger Monte Carlo uncertainty than the final 100-run comparisons. In particular, unavailable history matrices must not be interpreted as zero error or ignored when comparing that method.

Training and evaluation used the same post-calibration measurement-error condition. Transfer to a new error regime was not tested. Model selection used noisy validation references and may favor particular checkpoints; validation stopping does not establish global convergence. The tested input-dependent rule has more parameters than the learned fixed weights. The comparison therefore measures their final performance under the stated training procedure, rather than isolating parameter count, regularization, or sample efficiency.

Future work should obtain independent coordinate measurements with an explicit systematic-uncertainty budget, examine shared reference errors, and validate the sensor-error models using recordings. Comparing predictions based on longer histories and testing moving targets would determine the scope of the observed benefits. Those extensions should estimate position from observable inputs.

## Appendices

## Appendix A. Complete method comparisons

Table 4 reports every method evaluated in the final 100-run experiment. Training-covariance and learned-fixed results agree because the corresponding weights are algebraically identical for the shared training objective. Their agreement is an implementation check, not an independent replication. Single-input controls assess the benefit of combining the current measurement with the initial prediction. History methods assess the limits of compressing past measurements into an initial mean. All methods used the same evaluation targets and position measurements within each run. History-based GLS was available in all final runs.

**Table 4.**
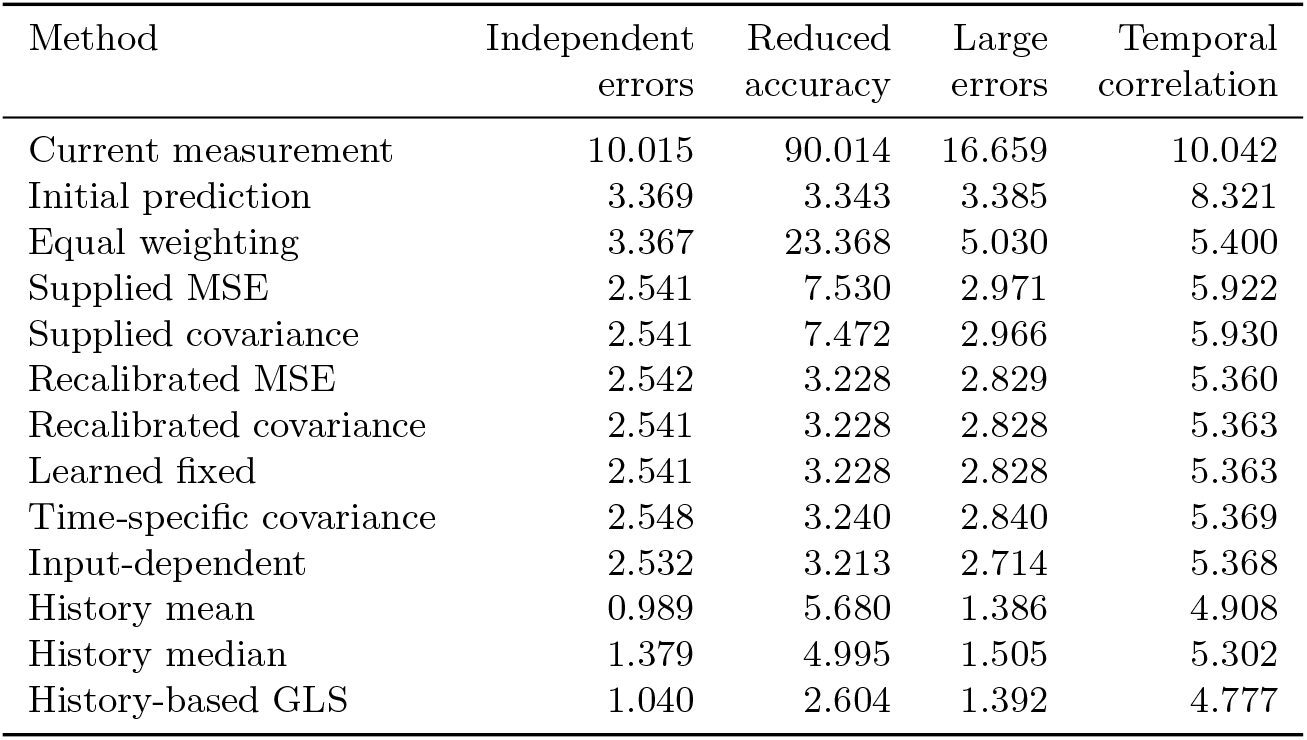
Complete final method results. Entries are mean run-level MSE in 10^*−*5^ m^2^, each based on 100 independently calibrated and trained runs. Each target remained fixed within its trial; all methods used observable position measurements. History methods retain more measurements after the first evaluated time point.

| Method | Independent errors | Reduced accuracy | Large errors | Temporal correlation |
| --- | --- | --- | --- | --- |
| Current measurement | 10.015 | 90.014 | 16.659 | 10.042 |
| Initial prediction | 3.369 | 3.343 | 3.385 | 8.321 |
| Equal weighting | 3.367 | 23.368 | 5.030 | 5.400 |
| Supplied MSE | 2.541 | 7.530 | 2.971 | 5.922 |
| Supplied covariance | 2.541 | 7.472 | 2.966 | 5.930 |
| Recalibrated MSE | 2.542 | 3.228 | 2.829 | 5.360 |
| Recalibrated covariance | 2.541 | 3.228 | 2.828 | 5.363 |
| Learned fixed | 2.541 | 3.228 | 2.828 | 5.363 |
| Time-specific covariance | 2.548 | 3.240 | 2.840 | 5.369 |
| Input-dependent | 2.532 | 3.213 | 2.714 | 5.368 |
| History mean | 0.989 | 5.680 | 1.386 | 4.908 |
| History median | 1.379 | 4.995 | 1.505 | 5.302 |
| History-based GLS | 1.040 | 2.604 | 1.392 | 4.777 |

As an additional analysis after completing the final comparisons, we aggregated the saved prediction errors separately at each evaluated time point (Fig. 3); no data generation, fitting, or parameter selection was repeated. History averaging improved as measurements accumulated under independent and large measurement errors, while the two-input methods retained roughly constant error in those conditions. Under temporal correlation, two-input errors also decreased as the separation between the initial and current measurements increased. Under reduced current accuracy, history averaging initially worsened as noisier measurements entered the mean, then improved as more measurements accumulated.

**Fig. 3.**
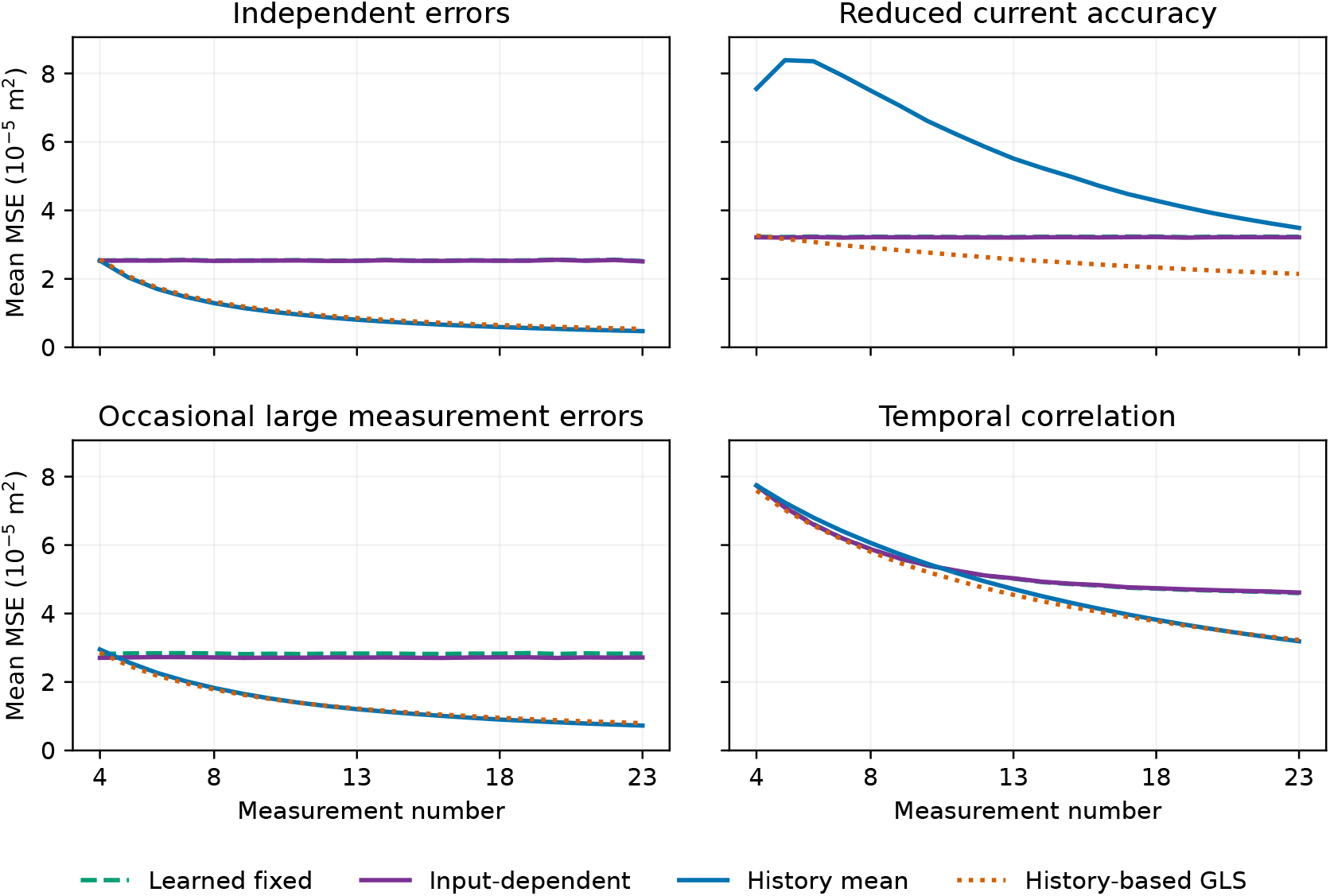
Time course of position-estimation error, computed by post-hoc aggregation of the frozen final predictions. Each curve averages coordinate-wise squared errors over 256 targets in each run and then over 100 independent complete runs. The first estimate uses the first measurement after the three initialization measurements; the last uses the final measurement of the trial. History methods use all available measurements, whereas the two-input methods combine the current measurement with the fixed initial mean. Curves show means without uncertainty bands; this display does not constitute an additional independent experiment.

At the first evaluated measurement, learned fixed weighting, input-dependent weighting, history averaging, and history-based GLS used the three initialization measurements and the current measurement. History averaging was equivalent to assigning weight 0.25 to the current measurement and 0.75 to the initial mean. Table 5 reports this additional aggregation of the saved predictions. In the large-error condition, input-dependent weighting had lower MSE than learned fixed weighting, history averaging, and history GLS in all 100 runs. This comparison removes the difference in the number of available measurements at this time point, but not differences in their representation or fitting procedures: GLS weights the four measurements individually, whereas the two-input methods compress the first three into their mean. This analysis used saved predictions without additional fitting or parameter selection.

**Table 5.**
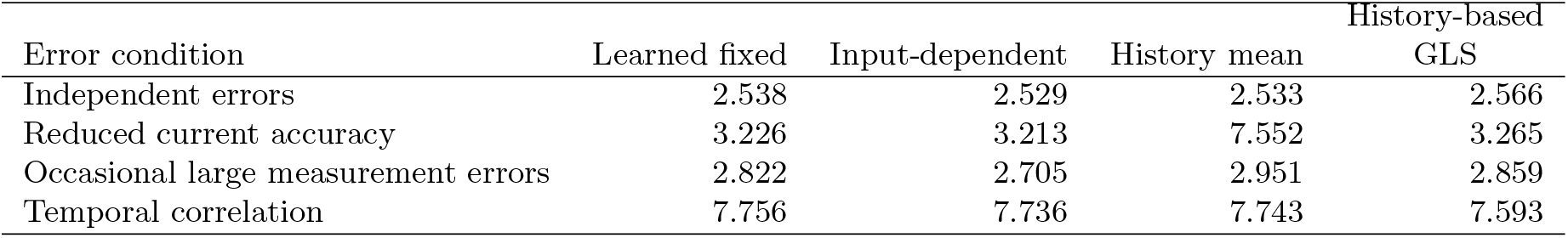
Error at the first measurement after the three initialization measurements. Entries are mean run-level MSE in 10^*−*5^ m^2^ over 100 complete independent runs per condition. All methods use the three initialization measurements and the current measurement; this table is a post-hoc aggregation of the frozen final predictions.

## Appendix B. Sensitivity to measurement assumptions

We examined nine measurement settings, including the primary setting, with 12 independent repetitions of each measurement-error condition. Each repetition used seven separate groups of 128 targets. All primary methods were retained. History-based GLS was unavailable when its noise-corrected training matrix was not positive definite: all 12 independent-error runs and seven large-error runs with fourfold reference-B SD. We did not regularize those matrices or replace the runs. GLS unavailability did not remove valid primary comparisons.

A coordinate standard with systematic offset *b* shifts corrected primary measurements and reference readings together. When other measurements are identical, translation correction yields a position shift *b* in every estimator, although the disagreement inputs and fitted weights can remain unchanged. Thus, learning from the references cannot identify the common coordinate error. With *e* denoting the original estimation error, the corresponding change in coordinate-averaged MSE is

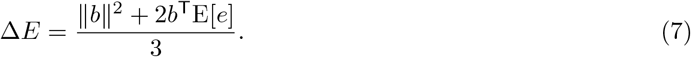

The experiment varies uncertainty about a coordinate reference; it does not let the estimator observe or correct the added standard bias.

Reference correlation hides shared error from the difference-based noise estimate. With equal marginal variance 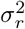 and correlation *ρ*, the mean reference has error variance 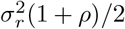, whereas the difference-based estimate approaches 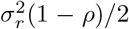. At *ρ* = 0.9, their ratio is 19. The true shared component was known only to the generator and evaluator, not supplied to the estimator. Unequal independent precisions alone do not invalidate the mean-reference correction: both expressions then give 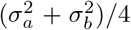. Finite translation-calibration residuals can still contribute biases and shared uncertainty.

Fig. 4 shows differences between learned fixed and input-dependent performance. Each cell uses independently generated measurements; it is not a paired contrast against the primary-setting cell. The large-error benefit was positive on average in every tested measurement setting, but its percentage reduction was smaller in the 5- and 10-mm standard-bias settings and the combined setting. Differences between settings include sampling variation and changes in the MSE denominator; they do not isolate an effect of bias on learning. Small differences in other measurement conditions changed sign. Twelve-run mean differences do not establish equivalent performance or a uniform benefit across reference errors.

**Fig. 4.**
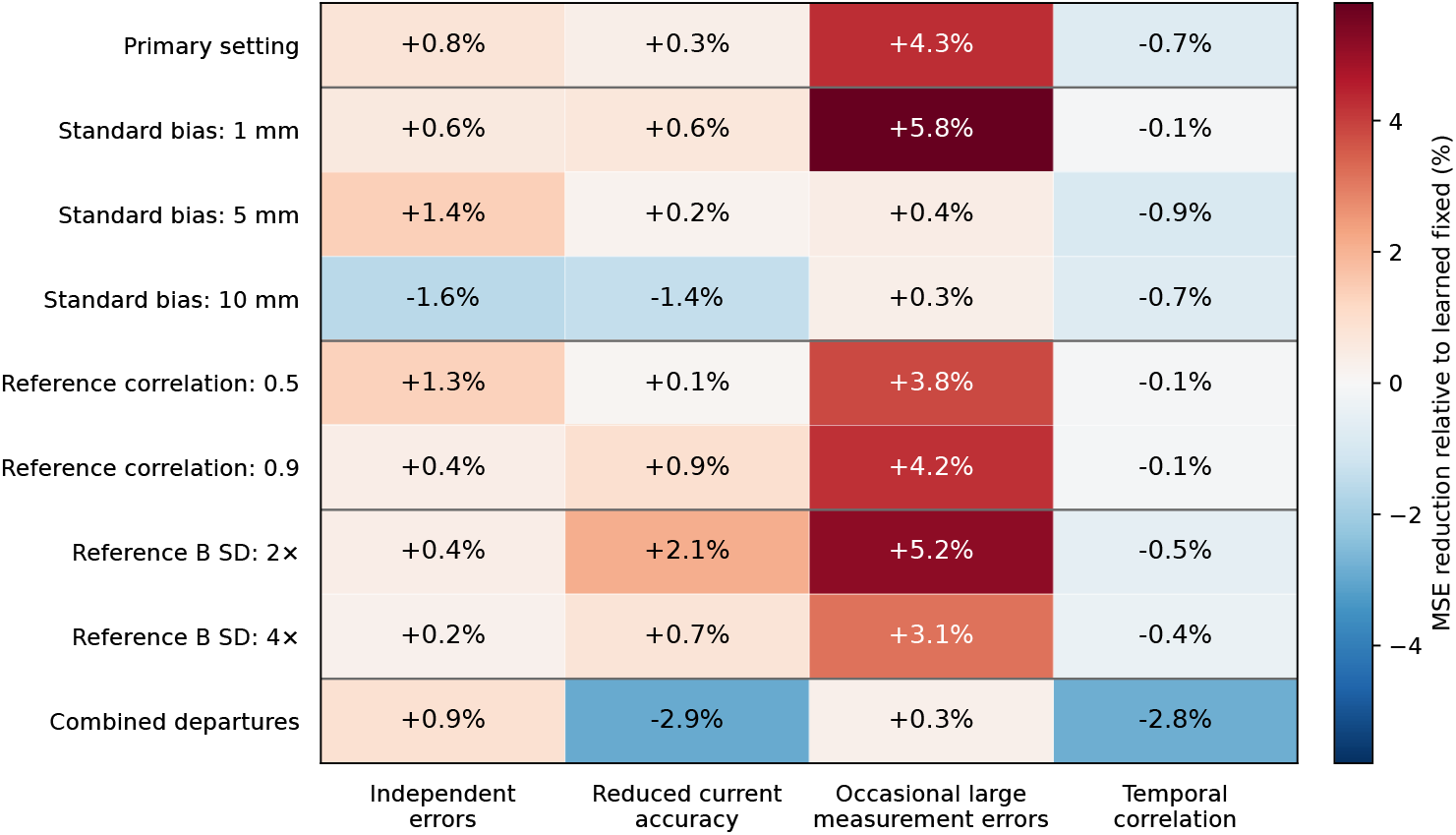
Development sensitivity of the input-dependent rule’s additional benefit. Each cell is the percentage MSE reduction relative to learned fixed weights, calculated from mean MSEs over 12 independent repetitions. Positive values favor the rule; negative values favor fixed weights. Standard bias is in one coordinate; reference correlations and SD factors are imposed generator parameters. The main 100-run experiment used a separate seed sequence and larger independent target groups.

## Statements and Declarations

### Ethics approval

Not applicable. This simulation study uses generated measurements and involves no human participants, animal experiments, clinical data, or recordings from real instruments.

### Consent to participate and consent for publication

Not applicable.

### Data availability

Generated measurements, fitted parameters, predictions, and analysis outputs for the revised study are available in version 0.2.0 of the Zenodo archive, 10.5281/zenodo.23117348. The standalone package is in `releases/observable-v0.2.0/` within the archived repository and includes SHA-256 manifests. Final and development results are distinguished; original processing of the same development measurements is preserved separately and is not an additional set of independent experiments. The original version 0.1.0, 10.5281/zenodo.20580788, preserves earlier materials excluded from the revised manuscript.

### Code availability

Revised simulation, verification, and analysis code is available under the MIT License in the same standalone package and in the GitHub version 0.2.0 release. Its README provides dependencies and reproduction instructions. The revised measurement simulation runs independently of the legacy musculoskeletal software retained elsewhere in the repository.

### Author contributions

Jun Kobayashi is the sole author and conducted the conceptualization, methodology, software development, experiments, analysis, visualization, and manuscript preparation.

